# Relation between the number of peaks and the number of reciprocal sign epistatic interactions

**DOI:** 10.1101/2022.01.23.477446

**Authors:** Raimundo Saona, Fyodor A. Kondrashov, Ksenia A. Khudiakova

## Abstract

Empirical essays of fitness landscapes suggest that they may be rugged, that is having multiple fitness peaks. Such fitness landscapes, those that have multiple peaks, necessarily have special local structures, called reciprocal sign epistasis ([14]). Here, we investigate the quantitative relationship between the number of fitness peaks and the number of reciprocal sign epistatic interactions. Previously it has been shown ([14]) that pairwise reciprocal sign epistasis is a necessary but not sufficient condition for the existence of multiple peaks. Applying discrete Morse theory, which to our knowledge has never been used in this context, we extend this result by giving the minimal number of reciprocal sign epistatic interactions required to create a given number of peaks.

## 1 Introduction

The fitness landscape is the relationship between genotypes and their fitness. Availability of high throughput methods and next generation sequencing started to experimentally characterize aspects of different fitness landscapes. Due to the enormity of the underlying genotype space ([12, 23]), the experimental approaches are limited to assaying fitness of: (a) closely related genotypes ([13, 16, 18, 19]); or (b), very restricted genotype spaces such as the interaction of a small number of protein sites ([11, 15, 22]). Nevertheless, the number of assayed genotypes in a single landscape is becoming larger in recent studies ([3, 17]) and it appears that the experimental characterization of a sufficiently large fitness landscape with multiple fitness peaks may be attainable within the next decade. Therefore, there is a need for development of computational methods ([1–3, 17, 22]) and theory ([24]) that can improve the description of experimental fitness landscape datasets, such as obtaining an estimate of the number of isolated peaks. Here, we use Morse theory to calculate the minimal number of reciprocal epistatic interactions for a given number of peaks on a landscape.

Epistasis is the interaction of allele states of the genotype, which shapes the fitness landscape. When the impact of allele states on fitness is independent of each other, there is no epistasis and the resulting fitness landscape is smooth and has a single peak. Epistasis can lead to a more rugged fitness landscape and decrease the number of paths of high fitness between genotypes. Epistasis that makes the impact of an allele state on fitness stronger or weaker is called *magnitude epistasis*. On the other hand, epistasis that causes the contribution of an allele state on fitness to change its sign (e.g., a beneficial mutation becomes deleterious) is called *sign epistasis* ([21]). When the two allele states at different loci change the sign of their respective contribution to fitness then this interaction is called reciprocal sign epistasis. In a simple example of this principle, in a two loci two allele model, there are four genotypes, 00, 01, 01 and 11. The following landscape is shaped by sign epistasis when genotypes 00, 01, 10 and 11 have fitnesses of 1,−1,1 and 1, respectively. Reciprocal sign epistasis is present when the fitnessses of 00, 01, 10 and 11 genotypes are 1,−1,−1 and 1, respectively.

Of course, the effect of an allele state can depend on more than just one other locus, or site, in the genome. When allele states in different loci impact each other then the epistasis is higher-order. Higher-order epistasis if found frequently in the characterized fitness landscapes ([20]), and it is clear that it has important evolutionary consequences ([4, 9, 10, 19]). However, models that allow studying such epistasis are at an early stage of their development (Crona et al. 2021 preprint, [5, 6]).

The evolutionary consequences of epistasis may be especially important when it leads to multiple local peaks. In that case, a population can get stuck on a suboptimal peak, decreasing the ability of evolution to find an optimal solution.

Using a combinatorial argument, [14] showed the following qualitative property: reciprocal sign epistasis is necessary for the existence of multiple peaks. In contrast, using Morse theory, we derive a more quantitative description of this relationship. This work might be the first formal use of Morse theory to study fitness landscapes.

## 2 Outline of the method

Morse theory studies the properties of some discrete structures (such as graphs) and special functions defined on them. In particular, the strong Morse inequality relates topological characteristics of a structure with the number of critical points of any function defined on it. Therefore, to use Morse Theory, we define a discrete structure that highlights reciprocal signed epistasis and a function based on the given fitness landscape.

The discrete strucuture is a graph: vertices are binary sequences (genotypes) and edges connect those genotypes within one-mutation distances. Moreover, we include edges between those vertices that are separated by reciprocal sign epistasis.

In the case of graphs, the only requirement for (Morse) functions is to assign a number to both nodes and edges. Naturally, the value on the vertices corresponds to the genotype’s fitness. On the other hand, the value on the edges is tailored for Theorem 1.

### Theorem 1

(Quantification of epistatic interactions) *Let genotypes be encoded as binary sequences. Consider a fitness landscape, that is a function that assigns a number to each genotype, with no strictly neutral mutations. Then*, # *reciprocal sign epistatic interactions ≥* # *peaks* − 1.

Because we model genotypes as binary sequences the sequence space is a hypercube. Also, we only consider fitness landscape with no strictly neutral mutation, i.e. all direct neighbours of a vertex must have a different value than this vertex. These two assumptions allows us to unambiguosly define reciprocal sign epistatic interactions.

## 3 Informal proof

Let us first briefly explain the combinatorial argument used in [14] to show the following qualitative property: reciprocal sign epistasis is a necessary condition for the existence of multiple peaks. The main idea is that between any two peaks (i.e. genotypes with locally maximal fitness) there must be a path consisting of single mutations connecting them. In particular, if the path is chosen well, the minimum fitness along this path is part of a reciprocal sign epistatic interaction. Theorem 1 is the corresponding quantitave version of this statement. In particular, if there are three peaks, we conclude that there is not only one reciprocal sign epistasis in the fitness landscape, but there must exist at least two of them.

Intuitevely, our result is explained by induction over the number of peaks as follows. The base case is when there are only two peaks, which is already explored in [14]. Then, for the inductive case, consider a fitness landscape and introduce a new peak on it. This new peak must be connected to all previous peaks through some reciprocal sign epistatic interaction. The question is if any of these interactions was not there berfore. We show that a new peak must introduce at least one more such interaction. To make this last step in the proof formal, we use discrete Morse theory.

## 4 Formal proof

The strong Morse inequality is a general tool that relates characteristics of a space with properties of special functions defined on it. In order to motivate subsequent definitions, let us present the original statement [8, Corollary 3.6, page 107] applied to graphs (instead of more general discrete structures).

### Theorem 2

(Strong Morse inequality) *Consider a graph G* = (*V, E*) *and a function f* : *V ∪ E →* ℝ. *Let b*_0_ *and b*_1_ *denote the first two Betti numbers of the graph G and let m*_0_*and m*_1_*denote the number of critical nodes and edges of f. Then, we have that*

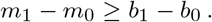

To use this result, we must define the following terms: Betti numbers and critical nodes and edges. But before we do that, note that if the number of reciprocal signed epistasis can be represented as a structural property of a graph (inside *b*_1_− *b*_0_), and the number of peaks can be encoded in a function (inside *m*_1_− *m*_0_), then this inequality allows us to quantify the necessary condition for the existence of multiple peaks.

We introduce all the necessary concepts before explaining the proof step by step.

### 4.1 Necessary definitions

In this section we introduce the terms used in Theorem 2 (*Betti numbers, critical nodes* and *critical edges*), as well as *reciprocal sign epistatic interactions*. All definitions coincide with those given in the general literature.

#### Definition 1

(Betti numbers) Let *G* = (*V, E*) be a graph. The zero-th Betti number (*b*_0_) is the number of connected components in *G*. The first Betti number (*b*_1_) equals |*E*| + *b*_0_− |*V* |, usually called cyclomatic number.

#### Remark 1

(Betti numbers in connected graphs) Let *G* = (*V, E*) be a connected graph. Then, *b*_0_= 1 and *b*_1_= |*E*| + 1 − |*V* |. Since *G* is connected, |*E*| *≥* |*V* | − 1, therefore *b*_1_ *≥* 0.

#### Definition 2

(Critical nodes and edges) Let *G* = (*V, E*) be a graph and *f* : *V ∪ E →* ℝ a function. We say that a vertex *v* ∈ *V* is critical if, for all edges *e* containing *v* we have that *f* (*e*) > *f* (*v*).

We say that an edge *e* = {*u, v*} ∈ *E* is critical if *f* (*e*) > max{*f* (*u*), *f* (*v*)}.

We denote *m*_0_ the number of critical vertices and *m*_1_ the number of critical edges.

#### Definition 3

(Reciprocal sign epistatic interaction) Consider a fitness landscape represented by *W* : 0, 1 _*n*_ R, where *n* is the length of the genotype. A reciprocal sign epistatic interaction is a collection of four different sequences *s*_1_, *s*_2_, *s*_3_, *s* _4_ ∈{0, 1}^*n*^ such that both sequences *s*_1_ and *s*_4_ are one single mutation away from *s*_2_ and *s*_3_ and it holds that

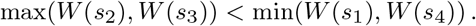

### 4.2 Proof

*Proof of* *Theorem* 1 Let a fitness landscape be respresented by a function *W* : {0.1}^*n*^ *→* ℝ, where *n* is the length of the genotype. Our proof consists in the following steps:

1. Define a graph.
2. Show that this graph is connected.
3. Define a function on the graph.
4. Apply the strong Morse inequality.

#### Definition of the graph

Consider a graph *G* = (*V, E*). Let *V* := {0, 1}^*d*^. Let set of edges *E* := *E*_1_ *∪ E*_2_ by defined in two steps: *E*_1_ and *E*_2_ has edges involving only sequences at Hamming distance one and two respectively. The set *E*_1_ contains only edges that connects a sequence with its fittest mutation, if it exists. Formally,

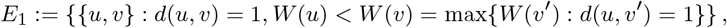

where *d* denotes the Hamming distance.

In the other hand, *E*_2_ contains edges that connect the two highest points of a reciprocal sign epistasis. Formally,

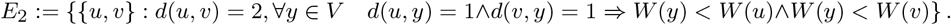

#### Connectedness of the graph

We now prove that *G* is connected, and therefore its first Betti number (*b*_0_, the number of connected components) is one. First note that any vertex is connected to a peak. Indeed, from any vertex, by following the path of fittest mutations, we can go to a peak by edges in *E*_1_. Therefore, we only need to prove that all peaks are connected.

By contradiction, assume that there are *K*_1_, …, *K*_*r*_ connected components of *G*. Note that in each component there might be multiple peaks. We will define a special path of single mutations. Formally, consider the “usual” sequence graph *G*_*S*_ = ({0, 1}^*d*^, 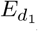), where 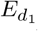 containing all edges connecting sequences at Hamming distance one. Take the path *P*_∗_ that connects two peaks in different components and has the highest minimum value, i.e.,

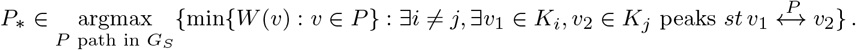

Without loss of generality, assume that *P*_∗_ connects *v*_1_ ∈ *K*_1_ and *v*_2_ ∈ *K*_2_. We will show that the two connected components *K*_1_ and *K*_2_ are in fact connected, which is a contradiction.

Denote *v*_*m*_ the vertex in *P*_∗_ that achieves the minimum fitness. Divide *P*_∗_ in by the path before and after *v*_*m*_, formally: 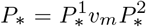. Our first observation is the following: all vertices in 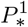 are in 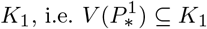, and similarly all vertices in 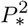 are in *K*_2_. Indeed, if it was not the case, consider 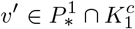. Since *v*′ ∉ *K*_1_, by following the fittest mutation, it is connected to a peak 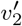 which is not in *K*_1_. Consider a new path 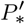 that goes from *v*_1_ to *v*′ and then to 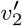. Note that the minimum fitness value in 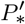 is higher than the one in *P*_∗_ and 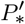 also connects two different connected components, which is a contradiction. Therefore, 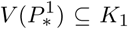. Similarly, we get that 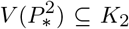. Having identified this property of *P*_∗_ and *v*_*m*_, we can construct a path in our graph of interest *G*, instead of *G*_*S*_.

Denote 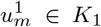 the vertex in 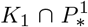 closest to *v*_*m*_, similarly denote 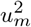 the vertex in 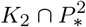 closest to *v*_*m*_. First notice that 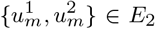, i.e. there a reciprocal sign epistasis between vertices with high fitness. Indeed, if this were not the case, we could connect them through another mutation that does not involve *v*_*m*_ and create a path 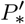 with a higher minimum value, which is a contradiction. Since 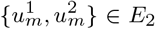, i.e. are connected in *G*, all we need to do to construct our desired path connnecting *K*_1_ and *K*_2_ is showing that 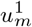 is connected to a peak in *K*_1_ and similarly 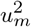 is connected to a peak in *K*_2_.

Since 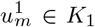, we can follow the fittest mutation path until a peak *u*_1_ ∈ *K*_1_ (similarly for 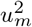 to a peak *u*_2_ ∈ *K*_2_). Consider the paths 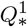 and 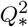, where 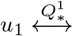 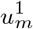 and 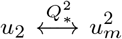. By definition of *E*_1_, we have that 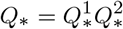 ⊆ *E*_1_. Finally, the path connecting two different peaks (assumed to be disconnected) is 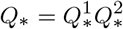.

Indeed, note that *Q*_∗_ is a path in *E* = *E*_1_ *∪ E*_2_ since 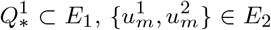 and 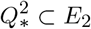. Therefore, the peaks *u*_1_ ∈ *K*_1_ and *u*_2_ ∈ *K*_2_ are connected. But this is a contradiction because *K*_1_ and *K*_2_ were two different connected components. This concludes the proof that *G* is connected.

#### Definition of a function

Consider the function *f* : *K →* R given by the following.

- For all *v* ∈ *V*,

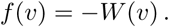
- For all *e* = *u, v* ∈ *E*_1_,

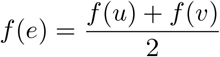
- For all *e* = *u, v* ∈ *E*_2_,

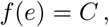

where *C* > max{−*W* (*v*) : *v* ∈ *V*}.

#### Application of Morse inequality

By Theorem 2, we have that

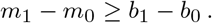

Since *M* is connected, we have that *b*_0_= 1. By definition of Betti numbers, and since *M* is connected, *b*_1_*≥* 0 (see Remark 1). The number of critical vertices is *m*_0_ and the number of critical edges is *m*_1_. By construction, the only critical vertices are peaks and the only critical edges are those in *E*_2_, i.e. edges that represent reciprocal sign epistasis. Therefore,

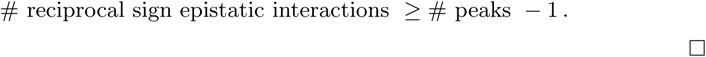

## 5 Discussion

We have shown that the multipeaked fitness landscape necessarily has no fewer pairwise reciprocal sign epistatic interactions than the number of fitness peaks minus one. This extends the result of [14] stating that the reciprocal sign epistasis is a necessary condition for multiple peaks. Additionally, our study showcases the application of discrete Morse theory to fitness landscapes.

As discussed in [14], reciprocal sign epistasis is not a sufficient condition for multiple peaks. Similarly, we do not show how to estimate the number of peaks from the number of epistatic interactions.

A sufficient condition for multiple peaks in terms of local interactions was given in a later work ([6]): reciprocal sign epistasis leads to multiple peaks if there is no sign epistasis in any other pair of loci.

The complication of deducing the global properties of fitness landscapes from the local properties of epistasis between specific sites arises due to the multidimensionality of the fitness landscape: local peaks formed by a pairwise epistatic interaction can be bypassed through a different dimension. Therefore, the condition formulated in terms of the pairwise epistatic interaction cannot be sufficient. One needs to know the full fitness landscape: to deduce that the fitness landscape has multiple peaks, one has to know that there is no sign epistasis in any other pairwise interaction ([6]).

For a quantitative result converse to ours, we anticipate that higher-order epistatic interactions have to be considered, which leads to the requirement of full information about the fitness landscape. We expect that this result can be obtained with a suitable definition of the higher-order epistasis. Such a result could be useful, for example, to study the empirical fitness landscapes if the number of mutations under consideration is small enough to make an almost complete description of the landscape feasible.

## Acknowledgments

We are grateful to Herbert Edelsbrunner and Jeferson Zapata for helpful discussions.

## Statements and Declarations

### Funding

Partially supported by the ERC Consolidator (771209—CharFL) and the FWF Austrian Science Fund (I5127-B) grants to FAK.

### Competing Interests

#### Declarations of interest

none.

### Data Availability

Data sharing not applicable to this article as no datasets were generated or analysed during the current study.

## Notes

### Competing Interest Statement

The authors have declared no competing interest.

## References

[1] Alley, E.C., Khimulya, G., Biswas, S., AlQuraishi, M. and Church, G.M., 2019. Unified rational protein engineering with sequence-based deep representation learning. Nature Methods, 16(12), pp.1315–1322. Available from: https://doi.org/10.1038/s41592-019-0598-1.

[2] Biswas, S., Khimulya, G., Alley, E.C., Esvelt, K.M. and Church, G.M., 2021. Low-N protein engineering with data-efficient deep learning. Nature Methods, 18(4), pp.389–396. Available from: https://doi.org/10.1038/s41592-021-01100-y.

[3] Bryant, D.H., Bashir, A., Sinai, S., Jain, N.K., Ogden, P.J., Riley, P.F., Church, G.M., Colwell, L.J. and Kelsic, E.D., 2021. Deep diversification of an AAV capsid protein by machine learning. Nature Biotechnology, 39(6), pp.691–696. Available from: https://doi.org/10.1038/s41587-020-00793-4.

[4] Canale, A.S., Cote-Hammarlof, P.A., Flynn, J.M. and Bolon, D.N., 2018. Evolutionary mechanisms studied through protein fitness landscapes. Current Opinion in Structural Biology, 48, pp.141–148. Available from: https://doi.org/10.1016/j.sbi.2018.01.001.

[5] Crona, K., 2020. Rank orders and signed interactions in evolutionary biology. eLife, 9, p.e51004. Available from: https://doi.org/10.7554/eLife.51004.

[6] Crona, K., Greene, D. and Barlow, M., 2013. The peaks and geometry of fitness landscapes. Journal of Theoretical Biology, 317, pp.1–10. Available from: https://doi.org/https://doi.org/10.1016/j.jtbi.2012.09.028.

[7] Crona, K., Krug, J. and Srivastava, M., 2021. Geometry of fitness landscapes: Peaks, shapes and universal positive epistasis. 2105.08469.9

[8] Forman, R., 1998. Morse theory for cell complexes. Advances in mathematics, 134(1), pp.90–145. Available from: https://doi.org/https://doi.org/10.1006/aima.1997.1650.

[9] Fragata, I., Blanckaert, A., Dias Louro, M.A., Liberles, D.A. and Bank, C., 2019. Evolution in the light of fitness landscape theory. Trends in Ecology and Evolution, 34(1), pp.69–82. Available from: https://doi.org/https://doi.org/10.1016/j.tree.2018.10.009.

[10] Kondrashov, F.A. and Kondrashov, A.S., 2001. Multidimensional epistasis and the disadvantage of sex. Proceedings of the National Academy of Sciences, 98(21), pp.12089–12092. https://www.pnas.org/content/98/21/12089.full.pdf, Available from: https://doi.org/10.1073/pnas.211214298.

[11] Kuo, S.T., Jahn, R.L., Cheng, Y.J., Chen, Y.L., Lee, Y.J., Hollfelder, F., Wen, J.D. and Chou, H.H.D., 2020. Global fitness landscapes of the shine-dalgarno sequence. Genome Research, 30(5), pp.711–723. Available from: https://doi.org/10.1101/gr.260182.119.

[12] Maynard Smith, J., 1970. Natural selection and the concept of a protein space. Nature, 225(5232), pp.563–564. Available from: https://doi.org/10.1038/225563a0.

[13] Melamed, D., Young, D.L., Gamble, C.E., Miller, C.R. and Fields, S., 2013. Deep mutational scanning of an RRM domain of the Saccharomyces cerevisiae poly(A)-binding protein. RNA, 19(11), pp.1537–1551. Available from: https://doi.org/10.1261/rna.040709.113.

[14] Poelwijk, F.J., Tănase-Nicola, S., Kiviet, D.J. and Tans, S.J., 2011. Reciprocal sign epistasis is a necessary condition for multi-peaked fitness landscapes. Journal of Theoretical Biology, 272(1), pp.141–144. Available from: https://doi.org/https://doi.org/10.1016/j.jtbi.2010.12.015.

[15] Pokusaeva, V.O., Usmanova, D.R., Putintseva, E.V., Espinar, L., Sarkisyan, K.S., Mishin, A.S., Bogatyreva, N.S., Ivankov, D.N., Akopyan, A.V., Avvakumov, S.Y., Povolotskaya, I.S., Filion, G.J., Carey, L.B. and Kondrashov, F.A., 2019. An experimental assay of the interactions of amino acids from orthologous sequences shaping a complex fitness landscape. PLOS Genetics, 15(4), pp.1–30. Available from: https://doi.org/10.1371/journal.pgen.1008079.

[16] Romero, P.A. and Arnold, F.H., 2009. Exploring protein fitness landscapes by directed evolution. Nature Reviews Molecular Cell Biology, 10(12), pp.866–876. Available from: https://doi.org/10.1038/nrm2805.10

[17] Russ, W.P., Figliuzzi, M., Stocker, C., Barrat-Charlaix, P., Socolich, M., Kast, P., Hilvert, D., Monasson, R., Cocco, S., Weigt, M. and Ranganathan, R., 2020. An evolution-based model for designing chorismate mutase enzymes. Science, 369(6502), pp.440–445. https://www.science.org/doi/pdf/10.1126/science.aba3304, Available from: https://doi.org/10.1126/science.aba3304.

[18] Sarkisyan, K.S., Bolotin, D.A., Meer, M.V., Usmanova, D.R., Mishin, A.S., Sharonov, G.V., Ivankov, D.N., Bozhanova, N.G., Baranov, M.S., Soylemez, O., Bogatyreva, N.S., Vlasov, P.K., Egorov, E.S., Logacheva, M.D., Kondrashov, A.S., Chudakov, D.M., Putintseva, E.V., Mamedov, I.Z., Tawfik, D.S., Lukyanov, K.A. and Kondrashov, F.A., 2016. Local fitness landscape of the green fluorescent protein. Nature, 533(7603), pp.397–401. Available from: https://doi.org/10.1038/nature17995.

[19] Visser, J.A.G. de and Krug, J., 2014. Empirical fitness landscapes and the predictability of evolution. Nature Reviews Genetics, 15(7), pp.480–490. Available from: https://doi.org/10.1038/nrg3744.

[20] Weinreich, D.M., Lan, Y., Wylie, C.S. and Heckendorn, R.B., 2013. Should evolutionary geneticists worry about higher-order epistasis? Current Opinion in Genetics and Development, 23(6), pp.700–707. Available from: https://doi.org/https://doi.org/10.1016/j.gde.2013.10.007.

[21] Weinreich, D.M., Watson, R.A. and Chao, L., 2005. Perspective: Sign epistasis and genetic constraint on evolutionary trajectories. Evolution, 59(6), pp.1165–74. Available from: https://doi.org/10.1554/04-272.

[22] Wittmann, B.J., Yue, Y. and Arnold, F.H., 2021. Informed training set design enables efficient machine learning-assisted directed protein evolution. Cell Systems, 12(11), pp.1026–1045. Available from: https://doi.org/https://doi.org/10.1016/j.cels.2021.07.008.

[23] Wright, S., 1932. The roles of mutation, inbreeding, crossbreeding and selection in evolution. Proceedings of the XI International Congress of Genetics, 8, pp.209–222.

[24] Zhou, J. and McCandlish, D.M., 2020. Minimum epistasis interpolation for sequence-function relationships. Nature Communications, 11(1). Available from: https://doi.org/10.1038/s41467-020-15512-5.

